# Neurobehavioural correlates of changing one’s mind in ADHD and OCD

**DOI:** 10.64898/2026.07.09.737533

**Authors:** Katharina Zühlsdorff, Jeffrey W Dalley, Trevor W Robbins, Sharon Morein-Zamir

## Abstract

Cognitive flexibility is an executive function that allows individuals to adjust behaviour in response to changing environmental demands. We assessed volitional switching under uncertainty, without rule-based learning, in the ‘Change Your Mind’ task. Nineteen patients with obsessive-compulsive disorder (OCD), 19 patients with attention-deficit hyperactivity disorder (ADHD) and matched control participants (20 per group) completed the task whilst undergoing a functional MRI scan. The task was a two-alternative forced choice paradigm where each stimulus was presented twice successively, with spurious feedback following the first presentation. This allowed participants the opportunity to repeat or change their response. Participants with ADHD changed their response more frequently than controls following a previously correct response, associated with reduced accuracy on the second trial. This was accompanied with smaller differences between change and repeat trials in the superior frontal gyrus, paracingulate gyrus and frontal pole compared to controls. Participants with OCD did not differ from healthy controls in their performance but exhibited greater activity on both change and repeat trials in the pre- and postcentral gyri than controls. These results point to distinct neurobehavioural differences in patients with ADHD and OCD underlying what is often termed more broadly inflexible behaviour.

## Introduction

Cognitive flexibility refers to the ease with which one can adjust mental processes to adapt behaviour to changing environmental demands (Dajani & Uddin, 2015). It is a central executive function, alongside other control processes such as inhibition and updating (Miyake et al., 2000). Higher levels of flexibility have been associated with positive life outcomes such as healthy ageing (Burke et al., 2019), creativity (Chen et al., 2014), and communication competence (Martin & Anderson, 1998). Conversely, impairments in cognitive flexibility may play a role in the pathophysiology of numerous neuropsychiatric disorders. For example, both patients with ADHD and with OCD exhibit impairments in behavioural flexibility (Boostra et al., 2005; Chamberlain et al., 2021).

Cognitive flexibility is a multidimensional construct, with behavioural paradigms assessing related but nonetheless differing aspects (Dajani & Uddin, 2015; Dias et al., 1996). Existing paradigms that assess flexibility, such as the Wisconsin Card-Sorting Task (WCST) (Grant & Berg, 1948), the CANTAB intra-dimensional/extra-dimensional set-shifting (IED) task (Chamberlain et al 2021) and probabilistic reversal learning (PRL) (Izquierdo et al., 2017), frequently capture reactive flexible responding, in which behavioural adjustments are triggered by external signals indicating a need to change. However, less is known about proactive flexibility, in which these adjustments are initiated by internally generated signals and volitional choices. In a previous study, we presented a novel behavioural paradigm that focuses on proactive flexibility (Zühlsdorff et al., 2023). Forty healthy participants completed a two-alternative forced choice task in which a masked stimulus was presented twice, with feedback orthogonal to their performance being presented after the first response. Following the second stimulus presentation, participants were free to either repeat or change their first response. This enabled us to assess the behavioural and neural correlates of ‘the road not taken’. Given the importance of proactive cognitive flexibility in everyday functioning, we sought to evaluate whether this process is altered in patient groups that have been reported to exhibit impairments in cognitive flexibility.

Attention-deficit hyperactivity disorder (ADHD) is a neurodevelopmental disorder affecting children, adolescents and adults and is characterised by hyperactivity, impulsivity and attention difficulties (Faraone et al., 2015). Individuals with ADHD show impairments across several executive functions, including cognitive flexibility, attention and inhibition (Nigg et al., 2005). Impairments in flexibility have been observed on set-shifting tasks such as the WCST, with higher perseverative and non-perseverative errors (Alaghband-Rad et al., 2021; Rohlf et al., 2012). Converging observations have been made on task-switching paradigms, showing that individuals with ADHD perform more slowly and make more errors following a switch (King et al., 2007; Rauch et al., 2012). However, recent contrasting computational evidence has reported that adults with ADHD show decreased reinforcement sensitivity and excessive choice switching on PRL pointing to more nuanced difficulties in flexibility (Aster et al., 2024). Additional evidence suggests that individuals with ADHD struggle to incorporate immediate feedback to update their learning (Gabay et al., 2018). Overall, these findings suggest that aspects of flexibility are impaired although the specific differences in behaviour may be task-dependent.

Aberrant neural processing relating to cognitive flexibility has also been reported. During task switching, adults with ADHD exhibited increased activation in the anterior cingulate cortex (ACC) and anterior insula (AI), whereas activation in areas involved in reinforcement learning such as the orbitofrontal cortex (OFC) and putamen was decreased (Berberat et al., 2022; Cubillo et al., 2010; Dibbets et al., 2010). PRL in ADHD groups has been associated with reduced activation of the left posterior parietal cortex and diminished ventral striatal signals associated with learning from the unchosen option (Aster et al., 2024). Similarly to the behavioural findings, the neural correlates of flexibility in ADHD are still not fully understood and may also be dependent on specific (executive function) task demands (Morein-Zamir et al., 2014).

As noted above, difficulties in cognitive flexibility are found in several neuropsychiatric conditions. Impaired cognitive flexibility is also a key feature of obsessive-compulsive disorder (OCD) (Robbins et al., 2019). OCD is a debilitating psychiatric condition characterised by intrusive thoughts or behavioural urges that can lead to checking, washing, worrying and anxiety (American Psychiatric Association, 2013). Patients with OCD show cognitive inflexibility on set-shifting tasks (Chamberlain et al., 2007; Vaghi et al., 2017). Slower tracking of the Trail Making Test Part B has also been reported, as well as increased perseverative errors on the WCST and worse performance (Shin et al., 2014; Snyder et al., 2015). Behavioural findings on probabilistic reversal learning (PRL) in OCD have been broadly consistent across studies, showing that individuals with OCD exhibit increased switching behaviour (Apergis-Schoute et al., 2024; Kanen et al., 2019; Remijnse et al., 2006).

Patients with OCD also exhibit changes in the neural circuits underlying flexibility. Functional MRI studies have found reduced fronto-striatal activation in OCD alongside perseverative errors in PRL tasks (Chamberlain et al., 2008; Remijnse et al., 2006). On task-switching paradigms, patients diagnosed with OCD show reduced activation of the OFC and dlPFC and increased activation of the putamen, ACC and postcentral gyrus (Chamberlain et al., 2008; Gu et al., 2008; Remijnse et al., 2013). Patients also show reduced fronto-striatal and striato-thalamic networks, linked to poorer performance on the Trails Making Task B (Kim et al., 2022). This largely converges with widespread under-activation across frontoparietal areas during shifting reported in OCD patients compared to controls, as well as reduced activation in the caudate and thalamus (Morein-Zamir et al., 2016; Vaghi, Vértes, et al., 2017). However, greater connectivity between the frontoparietal and default mode networks and increased activation in the superior parietal cortex has also been reported during task switching (Liu et al., 2023).

Despite the prominence of cognitive inflexibility in clinical presentation, the mechanisms underlying impaired flexibility in ADHD and OCD appear to diverge. For example, computational modelling reveals distinct profiles: following reversals participants with OCD exhibit heightened punishment learning rate, whereas those with ADHD show a reduced reward learning rate (Rodríguez-Herrera et al., 2025). In healthy control participants, connectivity between the right and left posterior parietal cortices predicted performance on this task, which was not observed in individuals with either OCD or ADHD. However, previous research has mostly focused on reactive rather than proactive flexibility. As discussed above, patients with ADHD and OCD generally show deficits across flexibility tasks, such as PRL, WCST and IED. Our previously published task (Zühlsdorff et al., 2023), which we tested in these two patient groups in the present study, has potential to distinguish ADHD from OCD.

In this study, we report the behavioural and neural correlates underlying the ‘Change Your Mind’ task in patients diagnosed with either OCD or ADHD compared with their sex- and age-matched control groups. In our previous study, we found that negative feedback and an incorrect first response independently resulted in increased changing behaviour (Zühlsdorff et al., 2023). Behaviourally, we hypothesised that individuals with ADHD would be more likely to change their response on the second trial following negative feedback due to a heightened sense of uncertainty compared with controls. For patients with OCD, we predicted reduced overall switching regardless of feedback, reflecting the need to alleviate uncertainty through repetitive actions. At a neural level, we anticipated increased activations in the ADHD group and reduced activity in the OCD group in the IFJ, AI, ACC and dlPFC, which are areas we have previously reported to be associated with changing a response (Zühlsdorff et al., 2023). The primary objective of this study is to unravel performance and the neural correlates of proactive cognitive flexibility in patients with ADHD and OCD.

## Methods

### Participants

Participants were 78 individuals, which were allocated to one of four groups: ADHD (N= 19, N females = 6, mean age = 27.7), ADHD control group (N = 20, N females = 6, mean age = 29.11), OCD (N=19, N females = 15, mean age = 35.15) and OCD control group (N=20, N females = 15, mean age = 37.79). Controls for the ADHD and OCD group were age- and sex-matched (Table 1). All volunteers were aged between 18 and 60 years (Mean=32.14, SD=9.98) and were recruited via the University of Cambridge Behavioural and Clinical Neuroscience Institute volunteer panel, using adverts in the Cambridge community or from the Cambridgeshire and Peterborough Foundation NHS Trust and from local support groups.

**Table 1.** Participant demographics and clinical characteristics.

| **Group** | **Mean age**<br>**(SD)** | **N females** | **Mean**<br>**verbal**<br>**IQ (SD)** | **OCIR**<br>**(SD)** | **Y-BOCS**<br>**(SD)** | **ASRS**<br>**(SD)** | **CAARS**<br>**(SD)** |
| --- | --- | --- | --- | --- | --- | --- | --- |
| Control<br>ADHD<br>(N=20) | 27.65<br>(5.83) | 6 | 115.9<br>(9.60) | 13.23<br>(6.88) | 0.75 (1.07) | 26.10<br>(7.20) |  |
| ADHD<br>(N=19) | 29.11<br>(7.72) | 6 | 115.6<br>(9.47) | 22.16<br>(11.20) | 1.42 (2.25) | 48.53<br>(11.80) | 49.3 (8.90) |
| Control OCD<br>(N=20) | 35.15<br>(11.6) | 15 | 117.5<br>(6.30) | 11.55<br>(6.80) | 0.55 (1.15) | 24.45<br>(8.24) |  |
| OCD (N=19) | 37.79<br>(10.1) | 14 | 113.3<br>(9.11) | 26.74<br>(13.65) | 18.95<br>(7.33) | 32.00<br>(9.97) |  |

Patients were diagnosed according to the DSM-IV criteria through an interview with a psychiatrist or clinical psychologist. OCD patients did not meet the DSM-IV criteria for any other Axis I disorders, except two who met the criteria for generalised anxiety disorder. Thirteen OCD patients were on serotonin reuptake inhibitors and one on a tricyclic antidepressant. Exclusion criteria for the OCD group were: substance abuse in the last three months, psychotic disorders, schizophrenia, bipolar disorder or ADHD. Nine of the ADHD patients were unmedicated (of which two were previously medicated), nine were prescribed methylphenidate and one was taking atomoxetine. On the day of testing, ADHD patients had not taken their medication for the last 24 hours. ADHD patients did not meet the DSM-IV criteria for any other disorders. Exclusion criteria for the ADHD group were: substance abuse in the last three months, psychotic disorders, schizophrenia, bipolar disorder or major depressive disorder or OCD.

Verbal IQ was assessed using the National Adult Reading Test score (NART) (Mean=115.56, SD=8.68) (Nelson, 1982). Additional clinical information was collected from the following questionnaires: Adult ADHD Self-Report Scale (ASRS), Conners’ Adult ADHD Rating Scales (CAARS), Obsessive Compulsive Inventory-Revised (OCI-R) and Yale-Brown Obsessive Compulsive Scale (Y-BOCS). All participants provided informed written consent and were reimbursed for their time and travel expenses, with ethical approval granted from the Cambridge Local Research Ethics Committee (08/H0308/65). These data were collected together with other data not reported here from the same participants (Morein-Zamir et al., 2014, 2016) and results from control participants were also previously reported (Zühlsdorff et al., 2023).

### Task Performance

The ‘Change Your Mind’ task has been described in detail previously (Zühlsdorff et al., 2023) and is also described in the **Supplementary Materials,** alongside details on neuroimaging acquisition. The primary outcome measure was the mean change in response on the second presentation of each trial (R2). A secondary outcome measure was R2 accuracy. Data were analysed using linear mixed-effects models (LMEs) implemented in R version 4.0.4 (R Core Team, 2021) using the **lme4** package (Bates et al., 2015). Fixed effects included feedback (positive vs negative), accuracy on the first response (R1: correct vs incorrect), group (control vs ADHD or control vs OCD), and all interactions. Post hoc pairwise comparisons of estimated marginal means were carried out using the **emmeans** package (Lenth et al., 2018; Pinheiro et al., 2021). Multiple comparisons were adjusted using Tukey’s method. Three different LMEs were assessed, differing in their random intercepts and interactions. Akaike Information Criterion (AIC), Bayesian Information Criterion (BIC), and log-likelihood were used to compare the models. The best-fitting model, which yielded the lowest AIC and BIC values, included random intercepts for subjects: Mean change R2 ∼ Feedback x R1 accuracy x Group + (1 + subject).

To examine the relationship between task performance and dimensional symptom severity, correlation analyses were conducted between mean changing behaviour and obsessive-compulsive symptoms (OCI-R total score), as well as ADHD symptoms (CAARS total score). Pearson’s correlation coefficients were calculated.

### Imaging data

#### First-level and higher-level models

Information on pre-processing can be found in the **Supplementary Materials.** A first-level fMRI linear model was fit using FEAT (FSL) (Woolrich et al., 2001) for each run and included the following event types: (1) R1 correct, (2) R1 incorrect, (3) positive feedback, (4) negative feedback, (5) R2 change and (6) R2 repeat. Equivalent event types for easy trials and six movement parameters (x, y, z, pitch, roll, yaw) resulting from the image realignment to control for movement artefacts were also included. Contrasts included only difficult trials and consisted of: (1) R1 incorrect vs correct, (2) negative vs positive feedback, and (3) R2 change vs repeat.

The second level model was constructed by averaging the first-level results for each participant across the three runs. The second-level contrasts were then entered into group-level (third-level) mixed-effects analyses using two-sample t-tests to compare the groups. Whole-brain results were thresholded with a cluster-forming threshold of Z = 3.1 and a cluster-corrected significance level of p < 0.05 to control for multiple comparisons (Woolrich et al., 2004). FSLeyes was used to create the figures (Smith et al., 2004). The main event types of interest were R2 change, R2 repeat, positive feedback, negative feedback, R2 change vs repeat and negative vs positive feedback.

#### Connectivity analysis

Additionally, whole-brain psychophysiological (PPI) interaction analyses were conducted in FSL. Three ROIs (‘seeds’) were selected based on regions reported to mediate proactive cognitive flexibility (Dajani & Uddin, 2015; Uddin, 2021). The regions selected were the AI, ACC and inferior frontal gyrus (IFG). A 5 mm sphere was placed in the centre of the respective region based on the Harvard-Oxford Atlas. Mean timeseries for each participant and each ROI were extracted.

This analysis assesses if there is an interaction between the ROI timeseries and a cognitive process which accounts for the neural responses observed in other brain regions (Neufang et al., 2008). This requires the addition of extracted time series, the response to the cognitive process of interest (change vs repeat) and an interaction term between them in the first-level model. Contrasts tested differences in connectivity in the change and the repeat condition in patients vs controls (ADHD vs controls or OCD vs controls). Two-sample t-tests for each seed region were conducted and clusters above a z-threshold of 2.5 and p<0.05 were considered significant.

## Results

### R2 change and accuracy in ADHD vs controls

A comparison of mean R2 change between ADHD and the matched control group revealed a significant effect of group (F(1,37)**=**7.85, p=0.008, η^2^p=0.17), with overall higher changing behaviour in the ADHD group (control: mean=0.28, ADHD: mean=0.38) (Fig. 1A). There was also a significant effect of feedback on R2 change across all participants (F(1,423)=129, p<0.001, η^2^p=0.23), with higher R2 change following negative feedback (negative feedback: mean=0.43, positive feedback: mean=0.23), however, there was no significant group x feedback interaction. R1 Accuracy also affected mean R2 change (F(1,423)=105, p<0.001, η^2^p=0.20). Mean change was higher when the first trial was incorrect (correct: mean=0.24, incorrect: mean=0.42). Furthermore, there was a significant Group x R1 Accuracy interaction (F(1,423)=4.08, p=0.044, η^2^p=9.55x10^-3^). *Post hoc* analyses found that there was higher switching in the ADHD group after a correct response compared to the control group (t(56)=3.40,p=0.007). There were no significant differences in mean changing after an incorrect R1.

**Figure 1.**
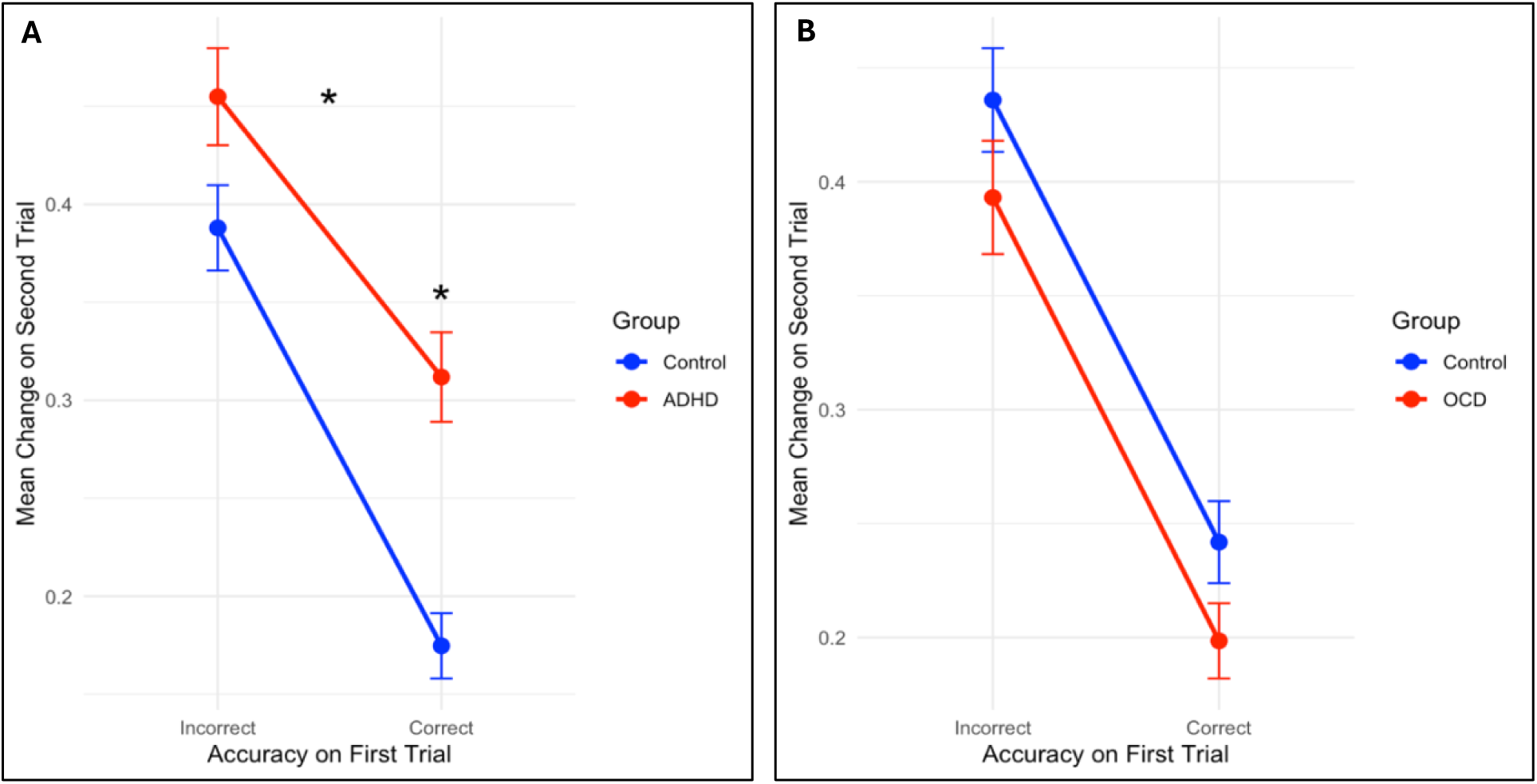
Effect of accuracy and group on R2 change. A) ADHD group: the linear mixed-effects model for R2 change indicated significant Group and Group x Accuracy on the first trial (Acc1) effects. Participants with ADHD changed on the second trial more overall, with an especially increased R2 change following a correct response on the first trial. B) OCD group: the linear mixed-effects model for mean R2 change showed no differences between the OCD and control groups. Error bars represent standard error of the mean. * - p<0.05.

As a secondary measure, we also assessed the mean accuracy on the second trial (mAcc2). There was a significant effect of group on mAcc2 (F(1,37)=6.22, p=0.017, η^2^p=0.14), with lower mAcc2 observed in the ADHD group (control: mean=0.60, ADHD: mean=0.55). R1 accuracy influenced the accuracy on the second trial (F(423)=271, p<.0001, η^2^p=0.39), with reduced accuracy following an incorrect first trial (correct: mean=0.74, incorrect: mean=0.41). We also found a Group x Acc1 interaction (F(1, 423)=26.76, p<.0001, η^2^p=0.06) (Fig. 2A). Post hoc comparisons showed that the accuracy on the second trial was lower in the ADHD group when participants were correct on the first trial (t(129)=5.39, p<.0001) but there was no group difference when participants were incorrect.

**Figure 2.**
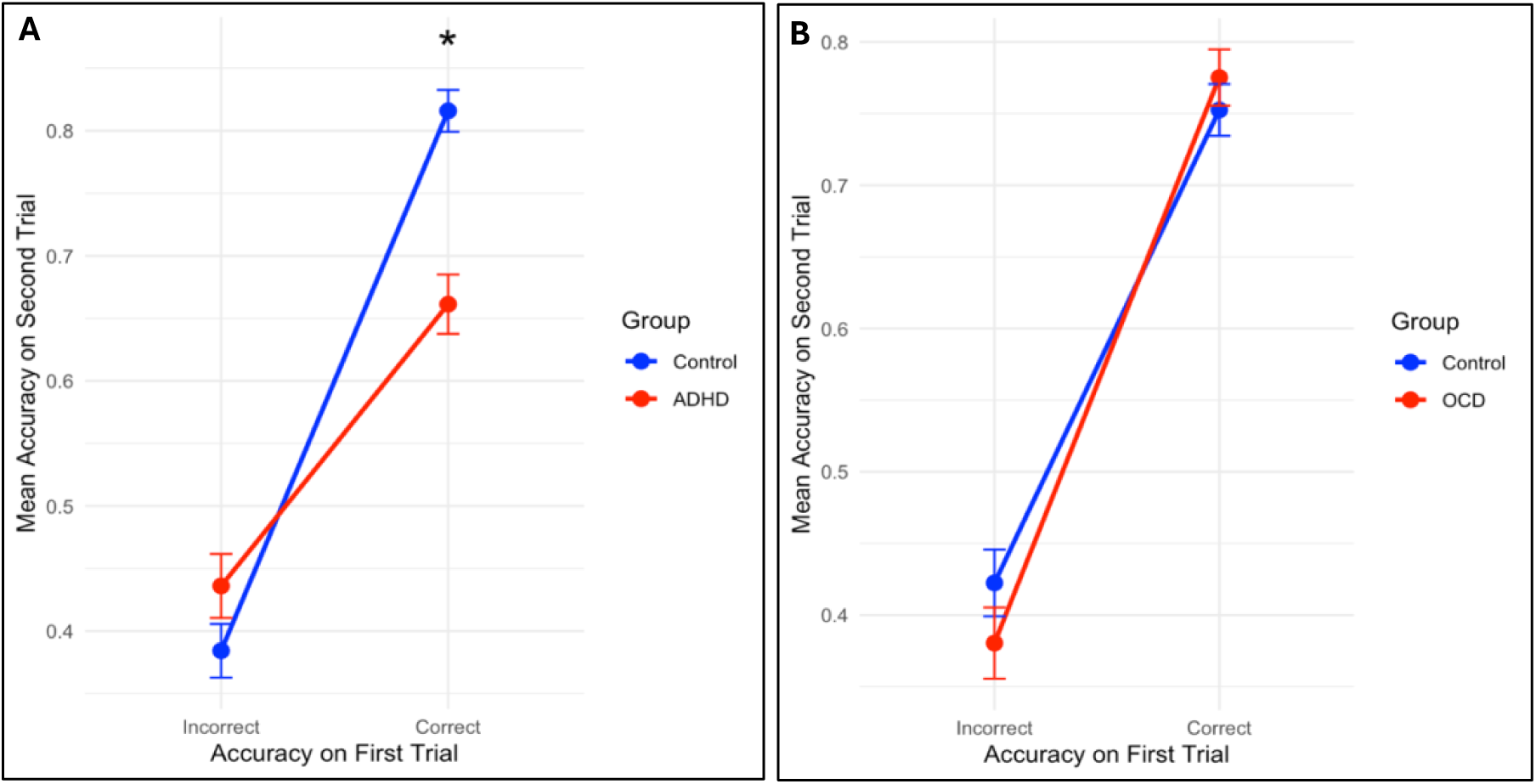
Effect of accuracy on the first trial and group on mean accuracy on the second trial. A) ADHD group: The linear mixed-effects model for mean accuracy on the second trial (mAcc2) had significant Group and Group x accuracy on the first trial (Acc1) effects.Participants with ADHD showed lower accuracy on the second trial when the first trial was correct. B) OCD group: The linear mixed-effects model for mAcc2 showed no differences between the OCD and control groups. Error bars represent standard error of the mean. * - p<0.05.

### R2 change and accuracy in OCD vs controls

Analyses of mean R2 change showed that feedback significantly affected mean R2 change (F(1,411)=42.86, p<0.001, η^2^p=0.09), with higher R2 change after negative feedback (positive: mean=0.27, negative: mean=0.37), however, there was no significant group x feedback interaction. R1 accuracy also affected mean R2 change (F(1,411)=152.41, p<0.001, η^2^p=0.27), with lower values following a correct response (correct: mean=0.22, incorrect: mean=0.42). However, no significant effects of group or interactive effects were found (Fig. 1B).

The secondary measure of mAcc2 also did not show any effects of group or any significant interactions (Fig 2B). There was a significant effect of Acc1 (F(1,447)=294, p<.0001, η^2^p=0.40). There was also a significant Feedback x Acc1 interaction (F(1,447)=21.19, p<.0001). Both positive feedback and correct Acc1 resulted in the highest mAcc2 compared to negative feedback and incorrect Acc1 (t(411)=12.3, p<.0001).

### Brain activations when changing versus repeating a response in ADHD vs controls

The primary imaging outcome was group differences in the activations underlying changing or repeating a response. There were no significant differences between the two groups in the change or repeat conditions separately. Comparing the change vs repeat contrast between the two groups revealed a larger difference between these two conditions in the control group than the ADHD group. This was observed in the superior frontal gyrus (SFG), paracingulate gyrus, right frontal pole and the right lateral occipital cortex (Fig. 3).

**Figure 3.**
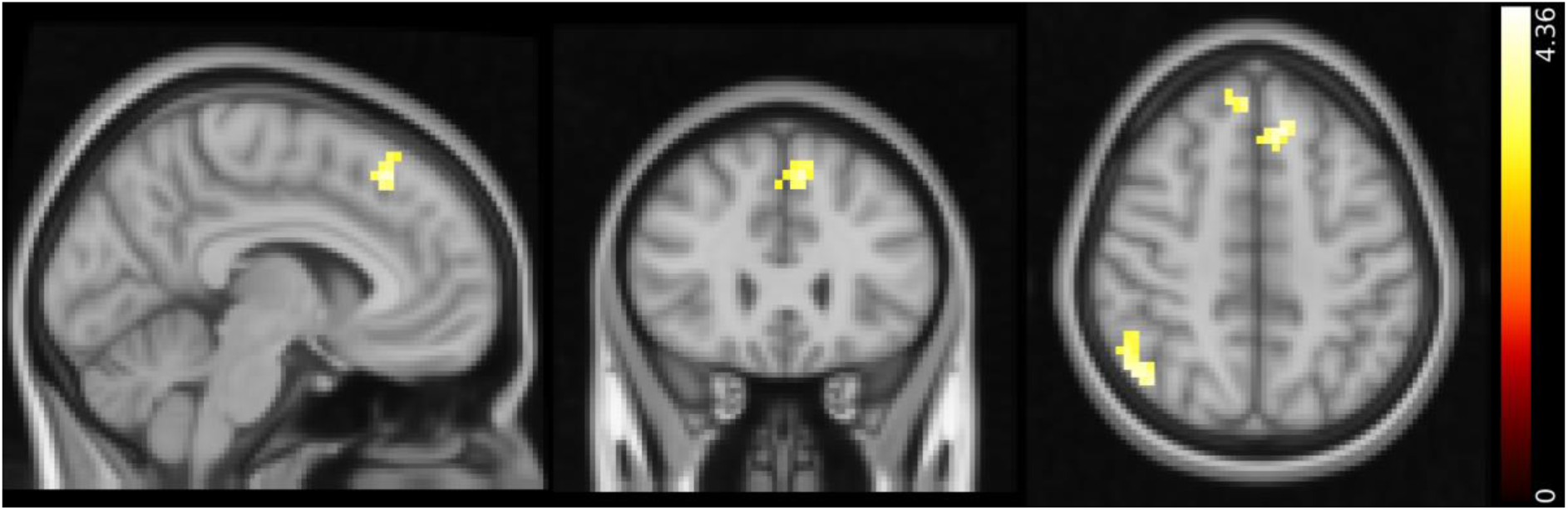
Results summary of the differences in change vs repeat contrast in control vs ADHD groups. Cross-section (MNI x=-6, y=27, z=48) of the contrast highlighting key regions activated more when participants changed compared to when they repeated their responses. This showed significantly greater differences between the two event types in controls than participants with ADHD, i.e. in controls, the difference in BOLD activity between the change and the repeat trials was greater than in individuals with ADHD. Activations detected with a whole-brain analysis involving one-sample t-tests with cluster thresholding with a Z-threshold of 3.1 and p<0.05. The colour bar represents the t-statistic.

### Brain activations when changing and repeating a response in OCD vs controls

When comparing the change and repeat contrasts in individuals with OCD vs controls, greater activity in the left pre- and postcentral gyri (including Brodmann area 4) in the OCD group were observed when both changing and repeating a response compared to control participants (Fig. 4). No differences in the repeat vs change contrast were observed between the two groups.

**Figure 4.**
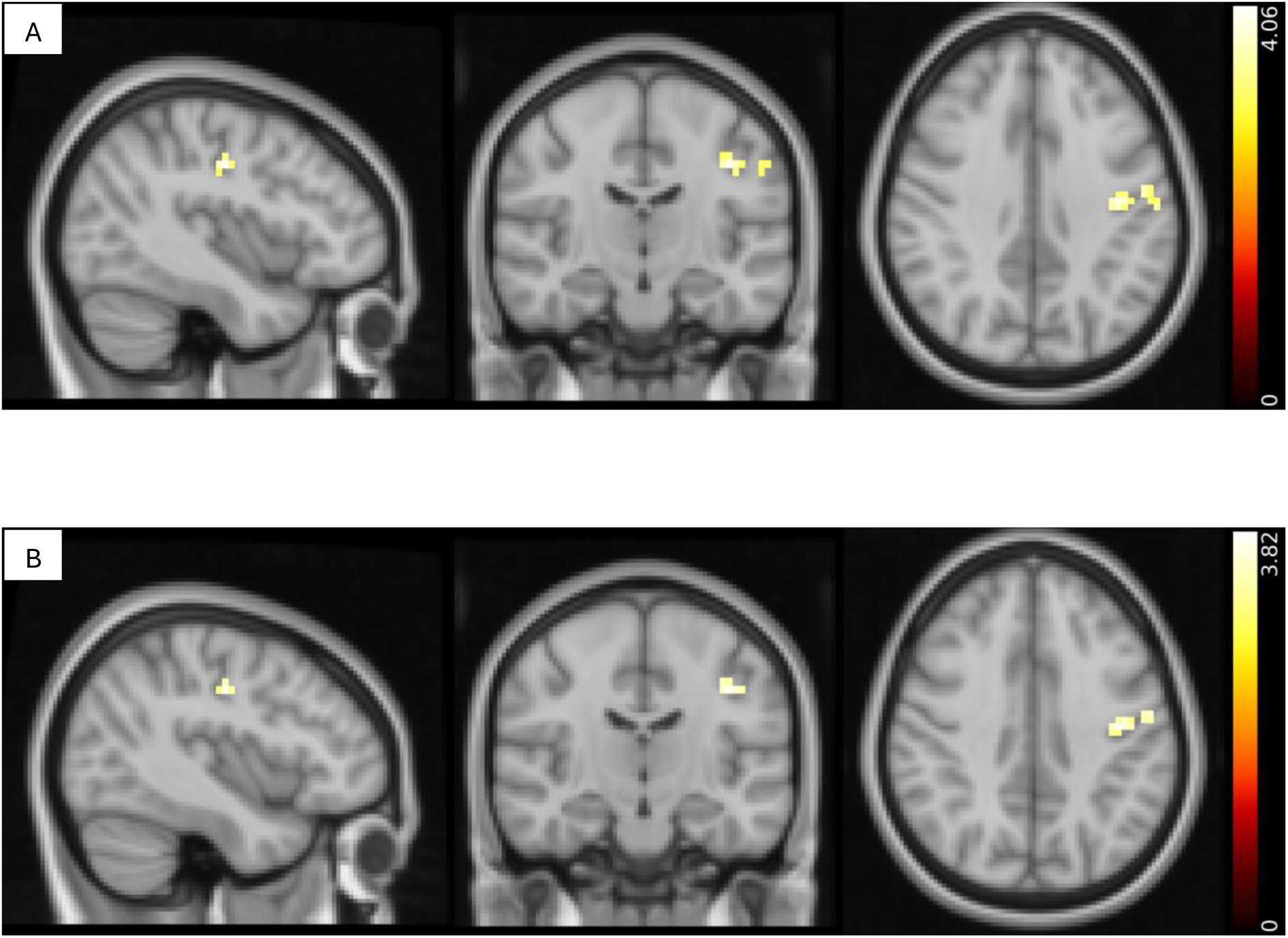
Results summary of the differences in the change and repeat contrast in control vs OCD groups. Cross-section of the contrast highlighting key regions activated more in the OCD group than in controls when participants A) changed (MNI x=-40, y=-16, z=38) or B) repeated (MNI x=-6, y=27, z=48) their responses (pre- and post-central gyrus) . Activations detected with a whole-brain analysis involving one-sample t-tests with cluster thresholding with a Z-threshold of 3.1 and p<0.05. The colour bar represents the t-statistic.

### Brain activations when receiving negative and positive feedback in patients

Furthermore, we investigated differences in brain activations following the presentation of positive or negative feedback. The contrast between the two feedback conditions was also compared between the two groups. When comparing the ADHD and control groups, no significant differences were found when positive or negative feedback were presented. However, the contrast between negative and positive feedback showed a greater difference in BOLD-dependent activity between these two events (negative vs positive feedback) in the control than in the ADHD group (Supplementary Figure 2). This was observed in the right IFG, right precentral gyrus and right lateral occipital cortex.

When receiving negative feedback, we observed that patients with OCD showed lower activity in the right and left SFG, cingulate gyrus and paracingulate gyrus compared to the control group (SF. 3). No significant differences during the presentation of positive feedback or in the negative vs positive feedback contrast were found.

### Neural connectivity when changing or repeating a response

The results from the PPI analyses in ADHD and their control group demonstrated differential interactions between brain regions when changing or repeating a response in the task. Specifically, we observed that during change trials there was lower connectivity between the IFG seed and the right and left precuneus cortex in the ADHD group (SF. 4). Additionally, during repeat trials there was greater connectivity in the same area and the right and left lingual gyri in the ADHD group compared to controls. No differences in the connectivity between the IFG and other brain regions were found in the change vs repeat contrast between the two groups. There were also no significant effects observed with the AI and ACC seeds.

A comparison of the OCD group with controls during change or repeat trials highlighted differences in functional connectivity between the AI and frontal pole in both change and repeat trials (SF. 5). Specifically, there was greater connectivity in the OCD group between the AI and left frontal pole during change trials, but reduced connectivity compared with the control group during repeat trials (SF. 6). No other significant effects where observed.

## Discussion

We report distinct behavioural and neuroimaging findings from ADHD and OCD patients performing the ‘Change Your Mind’ task. Our findings provide new information on an alternative form of flexibility and how it differs in these two patient groups. We replicated the findings from our previous study across all groups, specifically, that higher levels of changing are seen after negative compared to positive feedback (Zühlsdorff et al., 2023). Higher levels of changing were also seen when the response on the first trial was incorrect, which was also seen in our previous study. This form of flexibility has been poorly investigated to date: behavioural changes representing volitional control rather than task-set switching, rule-based learning or set shifting. Our results showed that patients with ADHD changed their responses on the second stimulus more frequently than controls, and even more so when their first response was correct, possibly indicating a lack of confidence in their first response. This tendency was associated with lower accuracy on the second trial following a correct response, i.e., they changed their mind more after their first response, despite their first response being correct. This behaviour was accompanied by a smaller difference in the haemodynamic BOLD response when contrasting change vs repeat trials in the SFG, paracingulate gyrus and frontal pole than in controls. On the other hand, patients with OCD, exhibited no behavioural differences on the task compared with the control group. There was increased activity in the left pre- and postcentral gyri in this group when both changing and repeating a response compared to the control group. They also showed reduced activity following presentation of negative feedback in the SFG, cingulate gyrus and paracingulate gyrus. Moreover, functional connectivity between the AI and frontal pole increased during change trials but decreased during repeat trials.

A prominent observation in this study was that individuals with ADHD changed their mind more often than controls and even when their first response was correct. This finding is consistent with previous behavioural evidence showing that ADHD is associated with increased switching on a PRL task and heightened trial-to-trial variability reflected in increased exploration rather than exploitation (Hauser et al., 2014; Luman et al., 2010). Studies investigating reinforcement learning mechanisms in ADHD have demonstrated that individuals show reduced sensitivity to positive feedback and have difficulties implementing successful response strategies (Frank et al., 2006; Groen et al., 2013). The increased switching we report in this study therefore likely reflect instability in employing goal-directed actions and an overreliance on reactive, feedback-driven responding. This may be due to an increased reliance on sensory feedback, rather than priors.

The ADHD group also showed a smaller difference between change and repeat trials in prefrontal regions in the ADHD group compared to control participants, possibly suggesting blunted recruitment of the frontoparietal network. One of the functions of this network is to support sustained attention (Kam et al., 2019; Prado et al., 2010). This points to impaired proactive control and potentially reduced confidence in internally generated decisions. The impaired proactive control in ADHD is reflected in lower thresholds to switch responses, resulting in less stable decision-making and is accompanied by a failure to recruit the appropriate neural networks required for switching, which include prefrontal cortical areas such as the SFG and paracingulate gyrus. There was also a smaller difference in the BOLD-response in the IFG during presentation of negative vs positive feedback in ADHD compared with controls. Previously, this same ADHD group showed more commission errors and reduced activation in the IFG during no-go trials on a combined shifting go/no-go task, suggestive of impaired response inhibitory control (Morein-Zamir et al., 2014). Other studies have also reported hypoactivation of the IFG and ventrolateral PFC, as well as overactivation of the visual cortex in patients with ADHD on flexibility tasks (Cortese et al., 2012; Rubia et al., 2005). The impaired control signal in the IFG previously noted in this sample as well as the differential response during feedback reported in this study may both contribute to the abnormal shifting behaviour exhibited in ADHD.

Consistent with the differences in clinical presentation between ADHD and OCD, the behavioural and neural findings in the OCD group were markedly different. This aligns with findings from the go/no-go task in the same samples (Morein-Zamir et al., 2014, 2016). Unexpectedly, task performance revealed no behavioural differences between the OCD and control groups, suggesting that flexible choice driven by volitional rather than rule-based control is unimpaired in OCD. The lack of behavioural differences contrasts with previous studies reporting impairments in OCD in several paradigms assessing flexibility, such as the WCST (Bohon et al., 2020), the IED set-shifting task (Chamberlain et al., 2021), deterministic reversal learning (Apergis-Schoute et al., 2024; Gu et al., 2008) and probabilistic reversal learning tasks (Remijnse et al., 2005; Apergis-Schoute et al 2024). Nevertheless, there is some occasional heterogeneity in performance of OCD patients on cognitive flexibility tasks (see Chamberlain et al 2021). Moreover, WCST performance was shown not to be impaired in adolescents with OCD (Marzuki et al 2021) and performance on the probabilistic reversal task in OCD is often characterised by reduced ’stickiness’ in response repetition (Apergis-Schoute et al., 2024; Kanen et al., 2019). Additionally, patients with OCD often do not show impaired flexibility in task switching paradigms (switch costs), instead showing overall slowing (Gu et al., 2008) or more errors (Liu et al., 2023; Remijnse et al., 2013). This pattern of findings is consistent with cognitive flexibility reflecting multiple component processes and OCD only being impaired in some (Dajani & Uddin, 2015; Gruner & Pittenger, 2017). In our previously published study on this OCD sample, which was free from comorbid depression, in a test combining Go/No-Go performance with set-shifting performance in the latter also did not differ from controls (Morein-Zamir et al., 2016). Moreover, two thirds of the patients were receiving SSRI medication which might have aided cognitive flexibility based on recent findings (Apergis-Schoute et al., 2017; Conceição et al., 2026).

Despite the lack of behavioural differences, at the neural level the OCD group exhibited some, albeit subtle, differences from controls. Greater activity in the left precentral and post central gyri was observed during both staying and changing in the OCD group compared to controls. This could be interpreted as evidence of greater cognitive effort to aid decisional performance. Altered connectivity between the frontal pole and AI specifically during change trials is consistent with prior findings (Grützmann et al., 2021; Stern et al., 2011) and potentially reflects heightened monitoring of decision outcomes supporting this compensatory interpretation. This change in connectivity may possibly be reflective of greater cognitive effort when changing behaviour, a process reliant on the interaction between the AI and the frontoparietal control network (Jiang et al., 2015). It is notable, however, that this same group of OCD patients, however, exhibited significant *under-*activation of the fronto-parietal network during set-shifting performance when behavioural performance was similarly preserved (Morein-Zamir et al., 2016). The SFG and cingulate cortex, which are involved in error monitoring (Carter et al., 1998), showed reduced activity following negative feedback in the OCD group. Although this contradicts previous studies showing increased activity in those areas (Fitzgerald et al., 2005; Stern et al., 2011), it could reflect generally altered feedback processing (Agam et al., 2014) and is consistent with a recent meta-analysis of task-based fMRI (Dzinalija et al., 2025).

Assessing ADHD and OCD using the change your mind task has allowed us to highlight different patterns of proactive flexibility despite the prominence of executive dysfunction and cognitive rigidity in the conceptualisation of both (Snyder et al., 2015). In the ADHD group, behavioural and neural abnormalities point to diminished top-down processing and impairments in proactive flexibility. In contrast, only subtle differences in the OCD were noted with intact performance and largely effective, sometimes apparently compensatory, recruitment of the relevant brain regions. When abnormalities were observed, they were associated with diminished neural responses to negative feedback and increased neural recruitment during the second response, regardless of whether participants repeated or changed their choice. As this task does not reliably distinguish between internal and external sources of uncertainty in changing one’s mind, it remains unclear how each source contributes to performance in the two patient groups. Disparate findings during current task performance are in line with the disparate clinical presentations of the two conditions, whereby executive dysfunction more broadly can contribute to impulsivity versus compulsivity (Faraone et al., 2015; Fineberg et al., 2014). A further limitation of the present study concerns differences in medication status across participants. Both the ADHD and OCD groups included individuals taking a range of medications, resulting in considerable heterogeneity within each clinical group. However, as the ADHD group did not take their medication prior to being tested, this group may be considered unmedicated. This was not the case for the OCD group. This variability may have influenced behavioural and neural measures and therefore makes it difficult to disentangle disorder-specific effects from those potentially attributable to pharmacological treatment.

In conclusion, this study highlights differences in volitional changing of behaviour in two patient groups characterised by cognitive inflexibility. Our findings suggest that this type of flexibility is selectively altered in patients with ADHD. In patients with OCD, we observed no behavioural differences on this particular type of flexibility but found subtle changes in neural processing that could be indicative of effective compensatory mechanisms. Our results do not highlight any decisional impairment or a lack of confidence in this group. These findings potentially inform future research on the neural and behavioural mechanisms of cognitive flexibility and guide the development of targeted interventions to improve adaptive behaviour in individuals with ADHD and OCD.

## Acknowledgments

We would like to thank Valerie Voon, Ulrich Muller, Jan van Niekerk, Tim van Hartefelt and Wolfgang Schwarzkopf for their assistance in collecting the data.

