## Supplementary Materials for "Neurobehavioural correlates of changing one’s mind in ADHD and OCD"

Supplementary Methods

**Task Procedure**

This task has been validated and previously published in (Zühlsdorff et al., 2023). Participants performed a two-alternative forced choice task and in which they were required to determine whether a target letter was ‘T’ or ‘L’ (see Fig. 1). Trials were presented in pairs, denoted by ‘1’ or ‘2’. The target letter was embedded in a row of Xs and masked after a brief delay. Participants were told that they would receive feedback after the first response, but that this feedback may not always be 100% accurate. Participants were also informed that after the feedback the same stimulus will be presented again and that they could change their mind on the second presentation. Participants were not informed that the feedback was orthogonal to performance and was monitored so that it was negative on half of accurate trials and positive on half of the incorrect trials.

All participants completed 56 pairs of trials in each of three runs, 48 of which were difficult and 8 were easy. Following a 500 msec fixation period, the target was briefly presented within a row of eight Xs in grey in the centre of the screen. After a brief period, the target was replaced with an ‘X’. The target location was counterbalanced, appearing in one of the 6 or 4 central locations on difficult or easy trials, respectively. On difficult trials, a staircase determined target duration, which was set to 60 msec initially and decreased by 10 ms following a correct response or increasing by 10 msec following an incorrect response. On easy trials, the target duration was set to 150 msec and was not adjusted. Stimuli were displayed for a total of 2 seconds. Participants had to press one of two buttons on a custom button-box. Response mappings on the button-box were counterbalanced across participants. Following 1 to 3 seconds (duration randomly selected at 100 msec intervals), feedback appeared for 1 second (‘+ correct +’ in green or ‘x wrong x’ in red). No monetary incentive was provided. After an additional interval (1-3 seconds), the second fixation appeared, followed by the same stimulus display and an intertrial interval (1-2.5 seconds). This procedure allowed for 6.2 seconds on average between the two responses of each pair. Before entering the scanner, participants had to complete 12 practice pairs. Trial order per block was randomised and the experiment was programmed in Visual Basic.

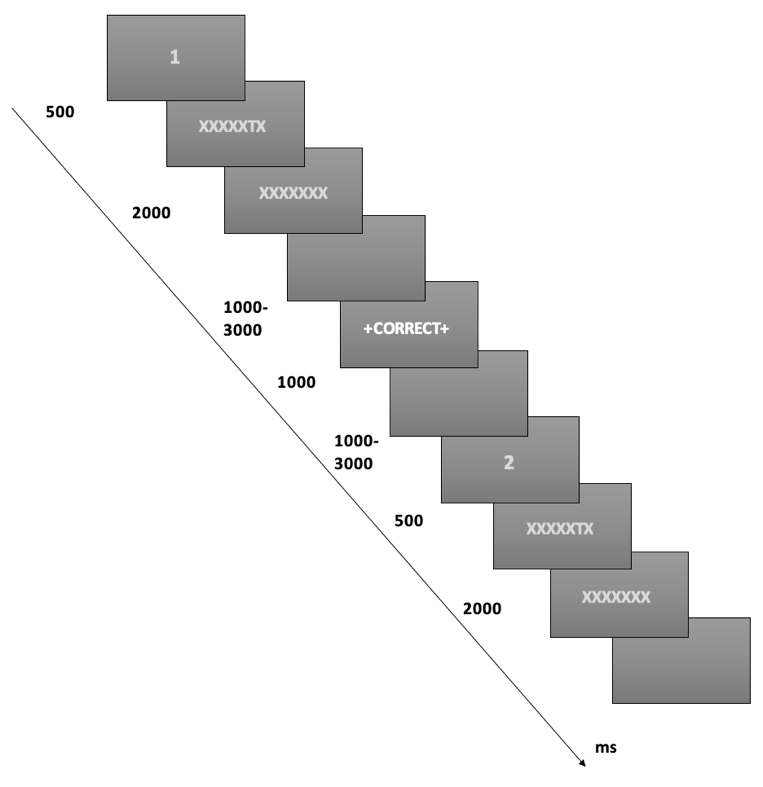

**Supplementary Figure 1.** **‘Change Your Mind’ task structure.** A target letter (‘T’ or ‘L’) was presented and masked with an ‘X’ after a predetermined duration in paired trials. The total stimulus duration of 2000 msec was comprised of a brief target presentation and a subsequent mask. Following their first response, feedback was presented that was orthogonal to their performance. This was followed by an identical second stimulus presentation. The first and second of each pair were denoted by fixation appearing as the numeral ‘1’ or ‘2’. Event durations are shown to the left of the task representation.

**Neuroimaging Acquisition**

Functional imaging was conducted on a 3 T Siemens MAGNETOM Trio scanner at the Wolfson Brain Imaging Centre (Cambridge, UK). A high-resolution T1-weighted anatomical image was acquired prior to functional runs. Functional images were collected using a standard Siemens echo-planar imaging (EPI) sequence sensitive to blood oxygenation level–dependent (BOLD) contrast (TR = 2000 ms, TE = 30 ms, flip angle = 78°) with an interleaved ascending acquisition order. The field of view was 192 × 192 mm with a 64 × 64 matrix, echo spacing of 0.47 ms, and bandwidth of 2442 Hz/Px. Each volume consisted of 32 axial slices (3 mm thickness; 3 × 3 mm in-plane resolution) aligned parallel to the anterior–posterior commissure line. Between 285 and 336 volumes were acquired per run.

**Imaging data**

*Pre-processing*

Preprocessing of the fMRI data was carried out using FMRIB’s Software Library (FSL) (Smith et al., 2004) and fMRIPrep, a Nipype-based pipeline (Esteban et al., 2018). Structural T1-weighted (T1w) images were corrected for intensity non-uniformity with N4BiasFieldCorrection and skull-stripped using antsBrainExtraction with the OASIS template from ANTs (Tustison et al., 2010). Nonlinear spatial normalization to the ICBM 152 Nonlinear Asymmetrical template (2009c) was performed with antsRegistration using brain-extracted versions of the T1w and template images (Avants et al., 2008). Tissue segmentation into cerebrospinal fluid (CSF), white matter (WM), and gray matter (GM) was performed on the skull-stripped T1w images using FSL FAST (Zhang et al., 2001).

Functional data underwent slice-timing correction (slicetimer, FSL) and motion correction (mcflirt, FSL) (Jenkinson et al., 2002). Susceptibility-induced distortions were corrected using field maps processed with fugue (FSL) (Jenkinson, 2003). Each functional run was then co-registered to its corresponding T1w image using boundary-based registration with six degrees of freedom (flirt, FSL) (Greve & Fischl, 2009). The distortion correction warp, BOLD-to-T1w transform, and T1w-to-template (MNI) warp were concatenated and applied in a single step using antsApplyTransforms with Lanczos interpolation.

For each run, framewise displacement was computed using the Nipype implementation (Power et al., 2014), and the first five volumes were discarded to mitigate T1 saturation effects. The preprocessed functional data were high-pass filtered (cut-off = 128 s) and spatially smoothed with a 6 mm full-width at half-maximum Gaussian kernel. A canonical hemodynamic response function was convolved with event onsets for subsequent modelling. Data quality was assessed by visual inspection of registration outputs, by confirming that no participant exceeded motion thresholds (defined as >10% of volumes flagged for excessive motion based on DVARS and framewise displacement), and by inspecting carpet plots. All participants met these criteria.

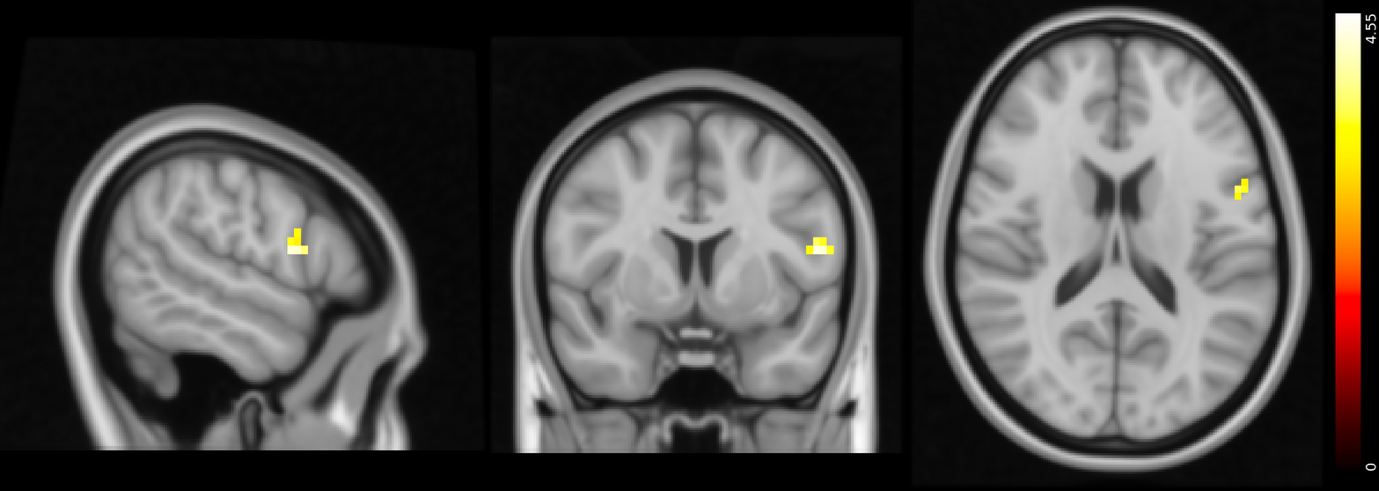
Supplementary Results

**Supplementary Figure 2. Negative vs positive feedback contrast in control vs ADHD groups.** Cross-section of the contrast highlighting key regions that showed a smaller difference in the negative vs positive feedback contrast in the ADHD group compared to control participants (MNI x=-54, y=7, z=16). Activations detected with a whole-brain analysis involving one-sample t-tests with cluster thresholding with a Z-threshold of 3.1 and p<0.05. The colour bar represents the t-statistic.

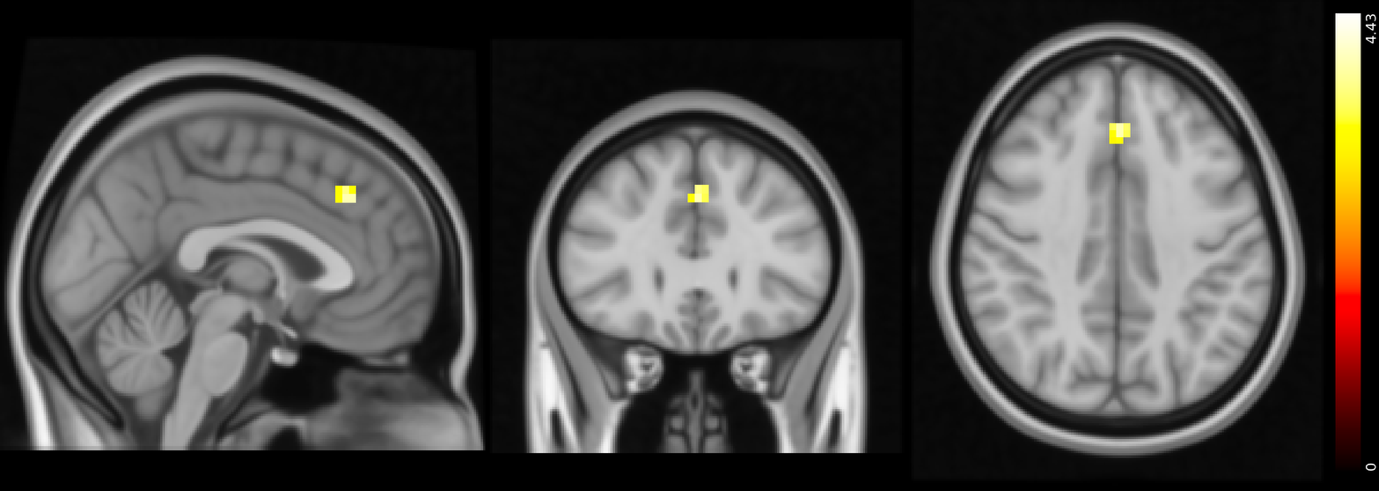

**Supplementary Figure 3. Results summary of the differences in the negative feedback contrast in control vs OCD groups.** Cross-section of the contrast highlighting key regions activated less in the OCD group than in controls when participants were presented with negative feedback (MNI x=-1, y=29, z=39). Activations detected with a whole-brain analysis involving one-sample t-tests with cluster thresholding with a Z-threshold of 3.1 and p<0.05. The colour bar represents the t-statistic.

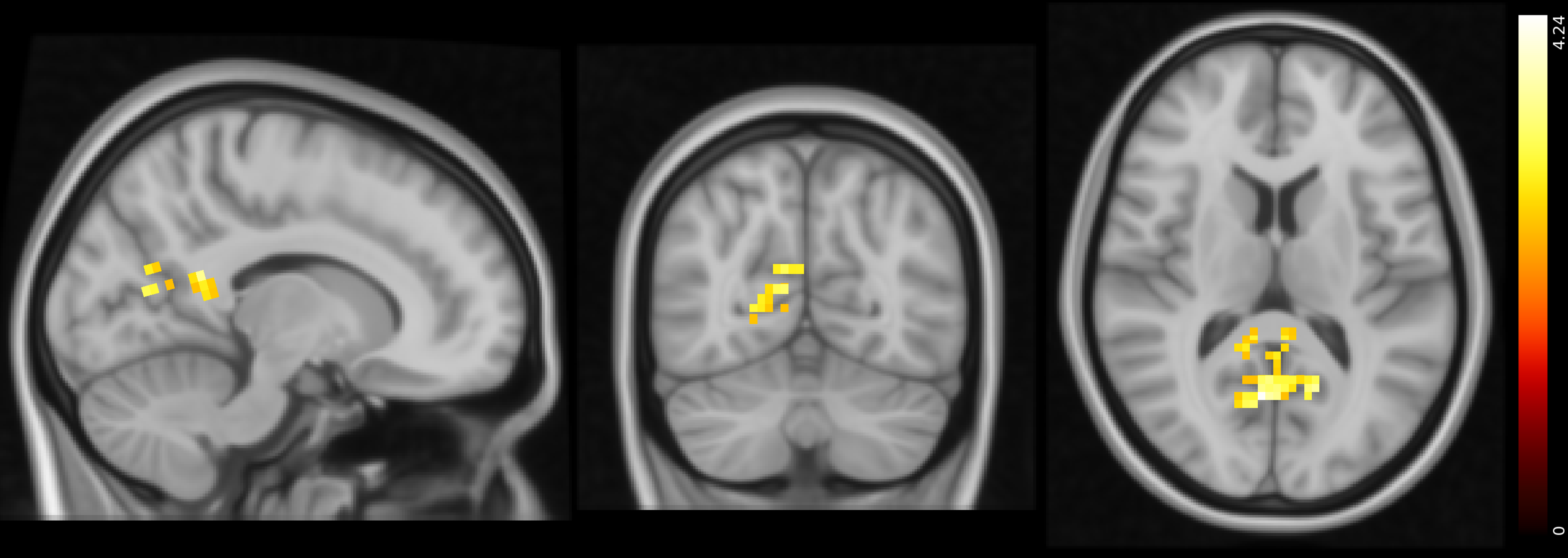

**Supplementary Figure 4. Results summary of the differences in PPI IFG analysis during change trials in control vs ADHD groups.** Cross-section of the contrast highlighting key regions that show lower connectivity with the IFG in the ADHD group than in controls when participants changed their response (MNI x=12, y=-65, z=12). Activations detected with a whole-brain analysis involving one-sample t-tests with cluster thresholding with a Z-threshold of 3.1 and p<0.05. The colour bar represents the t-statistic.

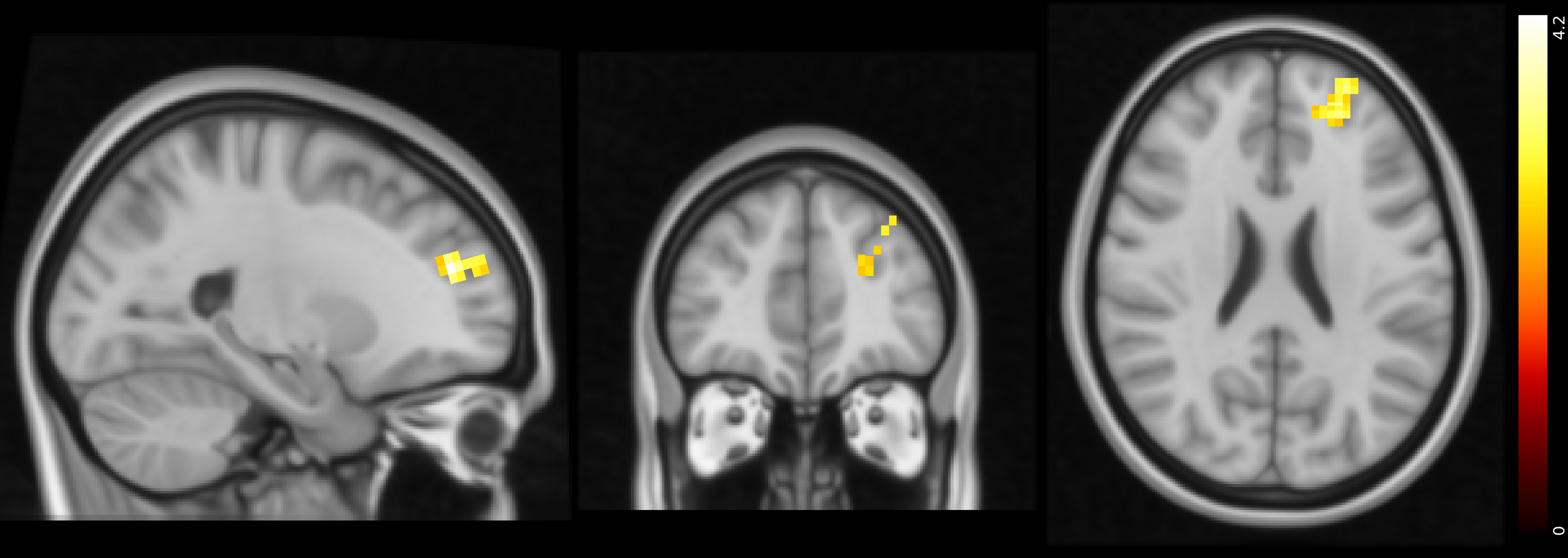

**Supplementary Figure 5. Results summary of the differences in PPI AI analysis during change trials in control vs OCD groups.** Cross-section of the contrast highlighting key regions that show greater connectivity with the AI in the OCD group than in controls when participants changed their response (MNI x=-23, y=43, z=23). Activations detected with a whole-brain analysis involving one-sample t-tests with cluster thresholding with a Z-threshold of 3.1 and p<0.05. The colour bar represents the t-statistic.

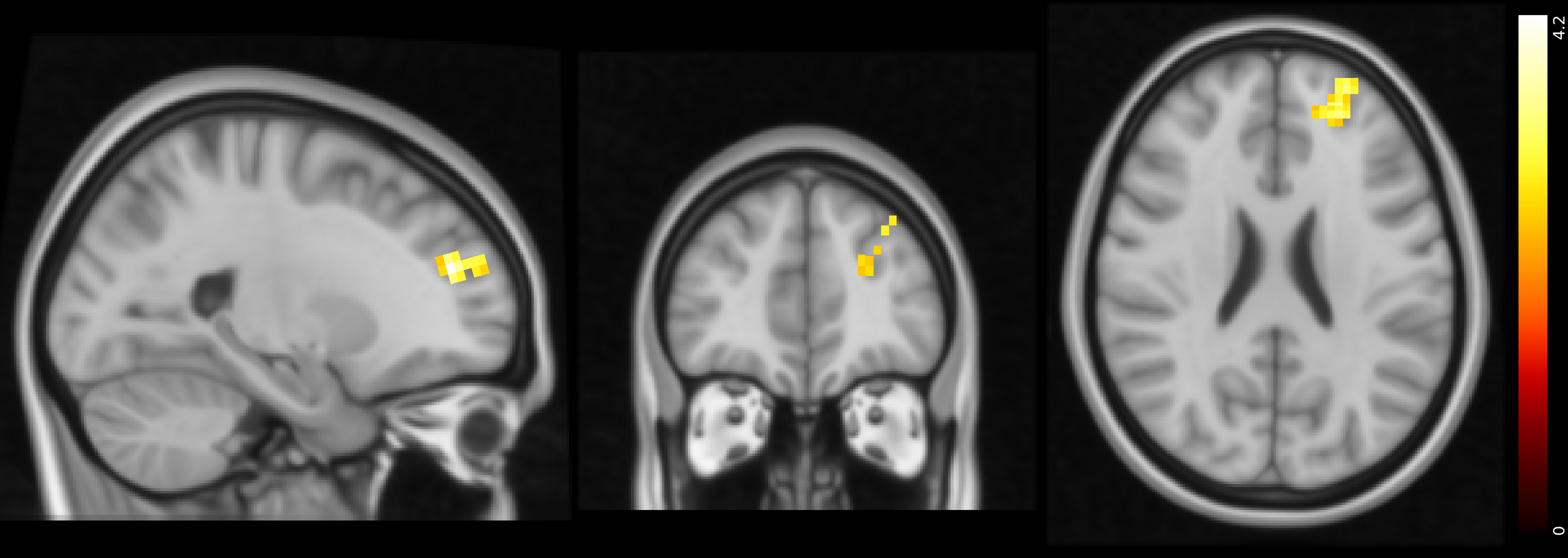

**Supplementary Figure 6. Results summary of the differences in PPI AI analysis during repeat trials in control vs OCD groups.** Cross-section of the contrast highlighting key regions that show lower connectivity with the AI in the OCD group than in controls when participants changed their response (MNI x=-23, y=43, z=23). Activations detected with a whole-brain analysis involving one-sample t-tests with cluster thresholding with a Z-threshold of 3.1 and p<0.05. The colour bar represents the t-statistic.
